# AutoPeptideML: A study on how to build more trustworthy peptide bioactivity predictors

**DOI:** 10.1101/2023.11.13.566825

**Authors:** Raul Fernandez-Diaz, Rodrigo Cossio-Pérez, Clement Agoni, Hoang Thanh Lam, Vanessa Lopez, Denis C. Shields

## Abstract

**Motivation:** Automated machine learning (AutoML) solutions can bridge the gap between new computational advances and their real-world applications by enabling experimental scientists to build their own custom models. We examine different steps in the development life-cycle of peptide bioactivity binary predictors and identify key steps where automation can not only result in a more accessible method, but also more robust and interpretable evaluation leading to more trustworthy models.

**Results:** We present a new automated method for drawing negative peptides that achieves better balance between specificity and generalisation than current alternatives. We study the effect of homology-based partitioning for generating the training and testing data subsets and demonstrate that model performance is overestimated when no such homology correction is used, which indicates that prior studies may have overestimated their performance when applied to new peptide sequences. We also conduct a systematic analysis of different protein language models as peptide representation methods and find that they can serve as better descriptors than a naive alternative, but that there is no significant difference across models with different sizes or algorithms. Finally, we demonstrate that an ensemble of optimised traditional machine learning algorithms can compete with more complex neural network models, while being more computationally efficient. We integrate these findings into AutoPeptideML, an easy-to-use AutoML tool to allow researchers without a computational background to build new predictive models for peptide bioactivity in a matter of minutes.

**Availability:** Source code, documentation, and data are available at https://github.com/IBM/AutoPeptideML and a dedicated webserver at http://peptide.ucd.ie/AutoPeptideML.

## Introduction

Peptides are short amino acid chains with 3 to 50 residues with a great variety of therapeutical properties. They have gained a lot of attention from the pharmaceutical and food industries, as their versatility makes them excellent candidates for drug or nutraceutical discovery (1). In this context, there is a growing demand for predictive models that can accelerate the discovery or design of peptides targeting new properties or bioactivities (2).

Novel developments in machine learning algorithms have offered new models for predicting protein structure (3) or different molecular properties (4). Despite these advancements, developing and evaluating new models is still an arduous process that requires both domain expertise and technical skills (2). Thus, most predictive models target broad and general applications, while solutions for more narrow use cases, like specific peptide bioactivities, remain underdeveloped (2).

Here, we propose building an automated machine learning (AutoML) system to automate the development of custom bioactivity predictors. There are several benefits that the introduction of such a system would provide. First, AutoML solutions can reduce the time required for model development from weeks to hours (5). Second, they help democratize machine learning by enabling researchers without a computational background to build effective models (5). Third, they help to ensure that best practices are followed, which can lead to increased trust on ML predictors within the field (6). Fourth, they can greatly simplify and accelerate current strategies that require not only strong computational skills, but also very tedious experimentation through extensive trial-and-error (2). Overall, an AutoML tool for building peptide bioactivity predictors will allow experimental researchers to seamlessly introduce advanced modelling techniques into their experimental workflows in a matter of hours (see Figure 1).

**Fig. 1.**
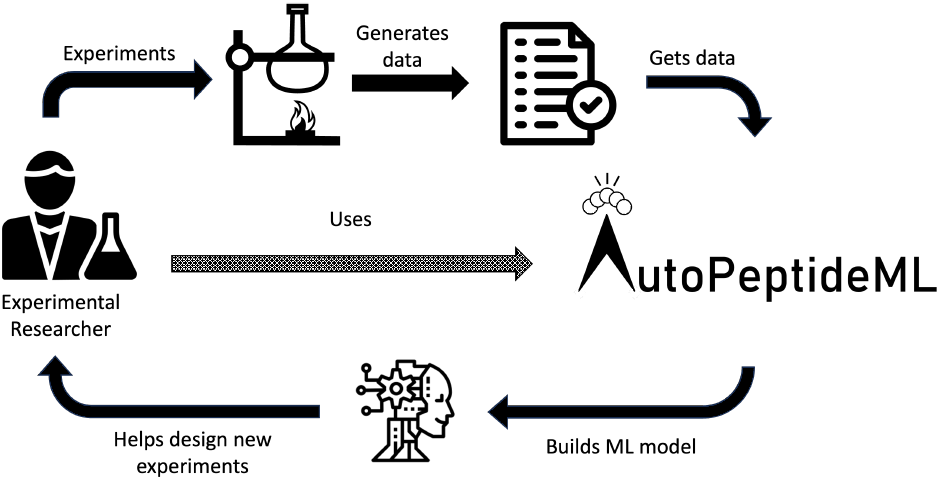
Integration of AutoPeptideML in an experimental workflow. AutoPeptideML allows experimental researchers to build new custom models from their data. These models can then be used to propose new experiments. The data generated from these experiments can, in turn, be used to generate a new improved model. This leads to a feedback loop where the experimentation is guided towards more relevant peptides with each iteration.

A review of the existing literature revealed five key steps in the development of a peptide bioactivity predictor that could be benefited by this automation: 1) data gathering for negative peptides, 2) dataset partitioning, 3) computational representation of the peptides, 4) model training and hyperparameter optimisation, and 5) reporting of model evaluation.

**1) Data gathering for the negative peptides**. Binary classifiers require both positive and negative examples. Finding positive examples for peptide bioactivity is relatively simple as there is prior literature describing the function and role of a multitude of peptides (7). However, there are few repositories enumerating peptides that do not present a certain function or property (2). Further, there is no consensus in the literature as to how negative peptides should be chosen: some works opt for choosing fragments of proteins (8; 9; 10), others look for actual peptides (8; 11; 12; 13), and yet others use peptides with a known bioactivity that is different from their target (8; 14; 15).

If we consider the first and second approaches, the model learns to differentiate between peptides with the target bioactivity and random sequences (either protein fragments or peptides). However, the problem is that the model can exploit multiple confounding factors that do not have a direct bearing on the specific bioactivity, but that are related to the differences between generally bioactive peptides and random sequences. In the third approach, the opposite is true, positive and negative peptides may be so similar to each other that the model will be biased towards specific differential features between both bioactivities, hindering its ability to generalise.

In this paper, we explored introducing an intermediate solution: to draw the negative peptides from a database with multiple bioactivities. This approach generates a distribution of negative peptides that is as unbiased as possible (by covering several distinct bioactivities) as to generalise adequately, but that is similar enough to the positive peptide distribution (by also being bioactive peptides) as to minimise confounding factors.

**2) Dataset partitioning**. To evaluate predictive models it is necessary to divide the data into at least three distinct subsets: training, validation, and testing. The independence between the training and testing subsets is essential to obtain a reliable estimation of the future model performance (16). To achieve this independence, community guidelines recommend building testing sets that do not share homologous sequences with the training set either by homology reduction (16) or homology partitioning (17; 18). Despite this, most of the peptide bioactivity predictors reviewed (8; 10; 11; 13; 14; 19; 20; 21; 22; 23; 24) do not introduce any correction for homology when partitioning their datasets and those that do (9; 15; 25; 26; 27; 28), use high thresholds (80-90% of sequence identity, Table SF), still allowing for similar sequences to be in different sets, which could lead to the overestimation of model performance due to data leakage. Here, we explored the effect of introducing homology-based dataset partitioning for building testing subsets more suited for evaluating model generalisation.

**3) Computational representation of the peptides**. For a predictive model to be able to interpret the peptide sequences, they need to be translated into mathematical objects (vectors or matrices). The reviewed literature offers different options for performing this transformation that include statistics of: residue composition (10; 15; 19), evolutionary profile (29), or physico-chemical properties (10; 11; 24; 27). The consensus that can be drawn from the variety of different descriptor combinations is that each predictive task will require a different set of descriptors. Finding the optimal combination is a crucial and intricate step in the modelling problem (2).

The advent of Protein Language Models (PLMs) like the ESM (evolutionary scale modelling) (30; 31) or RostLab (ProtBERT, Prot-T5-XL, or ProstT5) (32; 33) families has allowed for much simpler and richer protein representations. Given a sequence *s*, these models have learned the probability that a residue will appear in position *i* given the rest of the sequence {*s* − *r*_*i*_}, *P* (*r*_*i*_|{*s* − *r*_*i*_}). This probability is related to concept of conserved and unconserved positions that is often used when analysing multiple sequence alignments (3). The models are trained on a vast set of sequences from the UniRef (34) or BFD (35) databases which include not only protein sequences, but also peptides. Moreover, at least, two prior studies have demonstrated that they can be used for representing peptides outperforming traditional description strategies (36; 37). However, there are many PLMs varied both in terms of size and learning method and it is not clear which may be the optimal choice for computing peptide representations. In this paper, we continue this line of research by addressing two questions: does model size have an impact on how suitable their representations are for describing peptides? and is there any significant difference between different classes of models?

**4) Model training and hyperparameter optimisation**. There are many different algorithms for fitting predictive models to a binary classification task and choosing between them is an extended trial-and-error task. Here, we considered an alternative approach: to use standard tools in the AutoML domain for performing hyperparameter bayesian optimisation of simple machine learning models and ensembling them.

These contributions have been integrated into a computational tool and webserver, named AutoPeptideML, that allows any researcher to build their own custom models for any arbitrary peptide bioactivity they are interested in. The webserver requires only minutes to build a predictor and its use is as simple as uploading a dataset with positive examples. It provides an output summary that facilitates the interpretation of the reliability of the predictor generated and it has an additional window supporting the use of the generated models for predicting the bioactivity of any given set of peptides.

## Materials and methods

### Data acquisition

18 different peptide bioactivity datasets containing positive and negative samples were used to evaluate the effect of the different methods. These datasets were selected from a previous study, considering the use of the ESM2-8M PLM for general peptide bioactivity prediction (36). The datasets ranged in size from 200 to 20,000 peptides (see Table SF). Here, they are referred to as the “original” datasets.

### Dataset with new negative peptides

For each of the original datasets, a new version was constructed using the new definition of negative peptides, termed “NegSearch”. The negative peptides were drawn from a curated version of the Peptipedia database “APML-Peptipedia” comprised of 92,092 peptides representing 128 different activities (see Figure SA). To avoid introducing false negative peptides into the negative subset, all bioactivities that may overlap with the bioactivity of interest were excluded (see Table SB). To ensure that the negative peptides were drawn from a similar distribution to the positive peptides and thus minimise the number of confounding factors, for each dataset we calculated a histogram of the lengths of its peptides with bin size of 5. Then, for each bin in the histogram, we queried APML-Peptipedia for as many peptides as present in the bin, with lengths between its lower and upper bounds. If there were not enough peptides, the remaining peptides were drawn from the next bin.

### Dataset partitioning

Two different partitioning strategies were used to generate the training/testing subsets: A) random partitioning and B) CCPart (18), a novel homology-based partitioning algorithm which creates an independent testing set ensuring that there are no homologous sequences between training and testing. Briefly, the algorithm calculates pairwise alignments among all dataset sequences to form a pairwise similarity matrix. It then clusters these sequences based on the similarity matrix using the connected components algorithm. Lastly, it iteratively transfers the smallest clusters to the testing set. This process ensures that there are no sequences in the testing set with homologs in the training set. The datasets generated through this strategy are referred to as “NegSearch+HP”.

In both cases, A) randomly partitioned or B) homology partitioned, the training set is further subdivided into 10 folds for cross-validation. This second division relies on random stratified partitioning, to create 10 cross-validation folds.

### Pairwise sequence alignments

The pairwise sequence alignments were calculated using the MMSeqs2 software with prior k-mer prefiltering (38). We considered that two peptides were homologous if they had a sequence identity above 30% using the longest sequence as denominator.

### Peptide representations

In order to evaluate the PLM peptide representations, the following methods (30; 31; 32; 33) were evaluated: ESM2-8M, ESM2-35M, ESM2-150M, ESM2-650M, ESM1b, ProtBERT, Prot-T5-XL-UniRef50, ProstT5 (sequence mode), and one-hot encoding as a non-PLM-based baseline.

PLMs generate as output a matrix *M* with shape *n* × *e*, where *n* is the number of residues in the peptide and *e* is the model embedding size (in this study *e* ∈ [320, 1280], depending on the model). Each row in this matrix corresponds to a residue-level representation. We obtain a peptide-level representation *r* by averaging across all residues: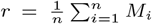. Please note that *r* is a vector with *e* dimensions.

### Model training and hyperparameter optimisation

In order to evaluate the model training and hyperparameter optimisation step, hyperparameter optimisation through bayesian optimisation (39) was performed separately for K-nearest neighbours (KNN), light gradient boosting machine (LightGBM) and random forest classifier (RFC) and all models were ensembled (see Table SC for more details about the hyperparameter optimisation). The optimisation aims to maximise model performance (measured as the average Matthew’s correlation coefficient (40) across the 10 cross-validation folds). The optimisation was conducted separately for each of the three models, leading to one optimal hyperparameter configuration per algorithm (three in total). After hyperparameter optimisation, each of the three models was trained against each of the 10 cross-validation folds using the optimal configuration. Thus, the final ensemble contained 10 instances (one per cross-validation fold) of each of the three models for a total of 3 × 10 = 30 models.

Final ensemble predictions were the average of all 30 individual predictions. This strategy is referred to as “Optimised ML ensemble” throughout the text. The three learning algorithms we used were chosen to provide a diverse representation of simple machine learning algorithms with computationally efficient implementations. We decided to use an ensemble, because it has been shown that for small datasets it leads to more robust predictors (41).

Our system was compared against an amended version of the UniDL4BioPep (36) framework, which we named “UniDL4BioPep-A” (more details about the architecture of the model can be found in the original publication (36) and are summarised in Table SD). This amendment differs from the original in that, following community guidelines (16), it used 10-fold cross-validation to determine the best possible checkpoint, instead of the hold-out testing set.

Every training experiment was run three times in order to get a crude estimation of the variability between experiment replications. The number of replicates is too small for proper statistical significance comparison, but the experimental design was constrained by the computational cost of each individual experiment run.

### Model evaluation metrics

Models performance was measured in terms of Matthew’s correlation coefficient (MCC), which is a binary classification metric that is specially recommended for measuring model performance in datasets with imbalanced labels (different number of positive and negative samples) (40; 42).

## Results and Discussion

We have focused our study of peptide bioactivity prediction in the binary classification task of discriminating between peptides that show a specific biologically-relevant property or function and those which do not.

### Automatic dataset construction

We first examined the effect of the two new methods proposed for automatically constructing the datasets: the sampling strategy for gathering negative peptides, and the homology-based partitioning algorithm (CCPart) used to generate training-testing subsets.

#### Effect of the sampling strategy for gathering negative peptides

Table 1 shows that the introduction of new negative peptides (column NS) has a limited effect on the interdependence between training and testing subsets going so far as increasing it in certain situations (when compared to the original datasets). On the other hand, Figure 2, shows that the change in negative peptides sampling strategy has a significant impact in apparent model performance with the most restrictive methods leading to the lowest apparent model performances. This result is consistent with the tumour T-cell antigen dataset experiencing an increase in model performance as the original set of negative peptides was comprised of non-tumoural T-cell antigens and is thus a more restrictive definition than generally bioactive peptides. Notably, the Antiviral dataset is the only case where performance increased after narrowing the negative class definition, and the reason behind this discrepancy remains unclear.

**Table 1.** Training-testing interdependence analysis of all datasets. Columns correspond to the different datasets constructed: Original datasets, the NegSearch datasets (NS) and the NegSearch datasets with homology-based partitioning (NS+HP). Values correspond to the proportion of training sequences with at least one homolog (sequence identity > 30%) in the testing set. *: Equivalent to Anticancer 1 for NS and NS+HP; **: Equivalent to Antimalarial 1 for NS and NS+HP

| Dataset | Original | NS | NS+HP |
| --- | --- | --- | --- |
| Antibacterial | 36% | 64% | 0% |
| ACE inhibitor | 1% | 3% | 0% |
| Anticancer 1 | 59% | 50% | 0% |
| Anticancer 2* | 30% | ~ | ~ |
| Antifungal | 34% | 58% | 0% |
| Antimalarial 1 | 41% | 15% | 0% |
| Antimalarial 2** | 10% | ~ | ~ |
| Antimicrobial | 47% | 63% | 0% |
| Antioxidant | 1% | 0% | 0% |
| Antiparasitic | 11% | 32% | 0% |
| Antiviral | 26% | 44% | 0% |
| Blood-brain barrier | 4% | 4% | 0% |
| DPPIV inhibitor | 5% | 4% | 0% |
| Anti-MRSA | 15% | 34% | 0% |
| Neuropeptide | 24% | 24% | 0% |
| Quorum sensing | 18% | 9% | 0% |
| Toxicity | 56% | 56% | 0% |
| Tumor T-cell antigens | 0% | 3% | 0% |

**Fig. 2.**
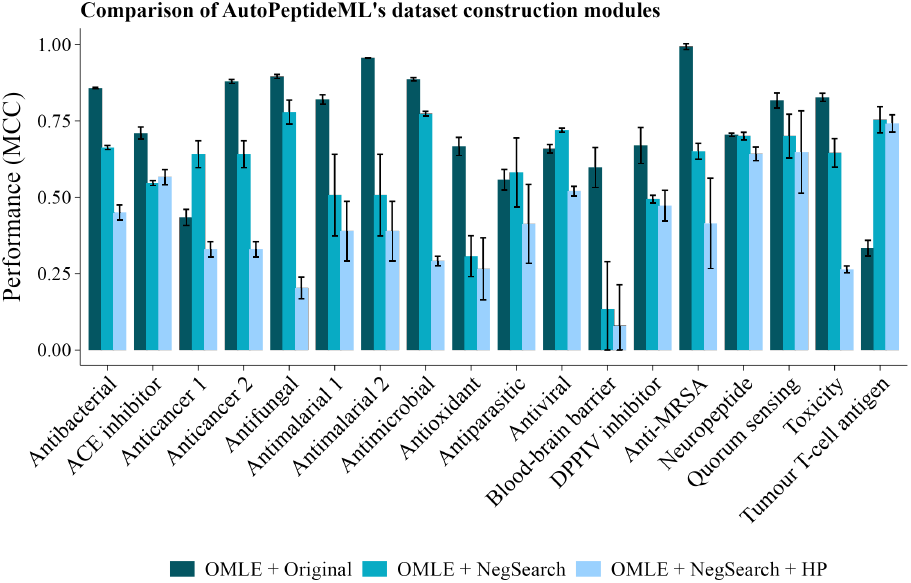
Evaluation of AutoPeptideML’s dataset construction modules. Error bars reflect the standard deviation across three replicates. OMLE: Optimised ML ensemble; Original: Original benchmark; NegSearch: Dataset with new negative peptides; HP: Homology-based dataset partitioning module.

It is particularly interesting to consider the change in the dataset pairs Anticancer 1 and 2, and Antimalarial 1 and 2 as the original datasets were built from the same sets of positive peptides, but relied on different sampling strategies for obtaining the negative peptides. Briefly, Anticancer 1 drew its negatives from a database of antimicrobial peptides (which are known to overlap with the anticancer peptides (43)) and Anticancer 2 drew them from a database of random protein fragments. This is reflected in Figure 2 where Anticancer 1 - Original demonstrates lower performance than Anticancer 2 - Original. It is noteworthy that the NegSearch dataset achieves an intermediate performance between them, as was to be expected from the NegSearch negative sampling being an intermediate definition between a specific bioactivity and random peptide sequences. Similarly, Antimalarial 1 - Original draws its negative peptides from a collection of random peptides, while Antimalarial 2 - Original draws them from a collection of protein fragments. Figure 2 shows how the more restrictive that the negative peptide sampling is the lower the model apparent performance, with Antimalarial 2 - Original achieving the highest apparent performance and Antimalarial - NegSearch the lowest.

Overall, the results indicate that the choice of sampling method for acquiring the negative peptides has an important effect on the perceived model performance. The optimal sampling method will depend in the inteded use for the model and whether it will be applied to protein fragments, random peptides or to distinguish between different peptide bioactivities. However, our experiments suggest that sampling from a collection with peptides with diverse bioactivities offers a balance between specificity and future generalisation, particularly, when the future target distribution is unknown at time of model development.

#### Effect of homology-based partitioning

We first analysed the interdependence between training and testing sets in the original datasets and compiled the results in Table 1 which indicates that for 13 of the 18 original datasets, at least 10% of the peptides in the training set are homologous to sequences in the testing set, compromising their independence. If we consider the datasets (see Table SF), for which homology-based independence correction was used (ACE inhibitor, Antioxidant, Antiparasitic, Anti-MRSA, and Neuropeptides) we see that in only two of them there is less than 10% of training sequences with at least one homolog in testing.

We also considered the effect of controlling the similarity between training and testing subsets by comparing model performance between the NegSearch datasets (where no homology correction was introduced) and NegSearch+HP (where we introduced homology-based partitioning). The effect observed across most datasets is a further drop in perceived model performance (Figure 2). The magnitude of the drop for some datasets suggests that prior studies have overestimated model performance and highlights the importance of introducing homology-based correction techniques when generating the training/testing subsets (16; 17; 18).

### Protein Language Models as peptide representation methods

Recent studies have reported the use of PLMs for predicting peptide bioactivity (36; 37), however, they have not been compared to a naive baseline representation like one-hot encoding; nor has there been an evaluation on which PLM may be more suited for peptide representation. All experiments are conducted with the NegSearch+HP datasets to ensure that we were properly evaluating model generalisation.

#### Baseline

First, we compare the PLMs to a naive baseline representation (one-hot encoding). Figure 3 shows that generally PLMs are significantly better representation methods across datasets, though in specific cases one-hot encoding appears to achieve similar performance. There are different idiosyncrasies within those datasets that may explain the behaviour, for example in the Blood-brain barrier experiments the number of training peptides is really small (∼200, see Table SF) which leads to a lot of instability between the different runs (as can be seen by the size of the error bars).

**Fig. 3.**
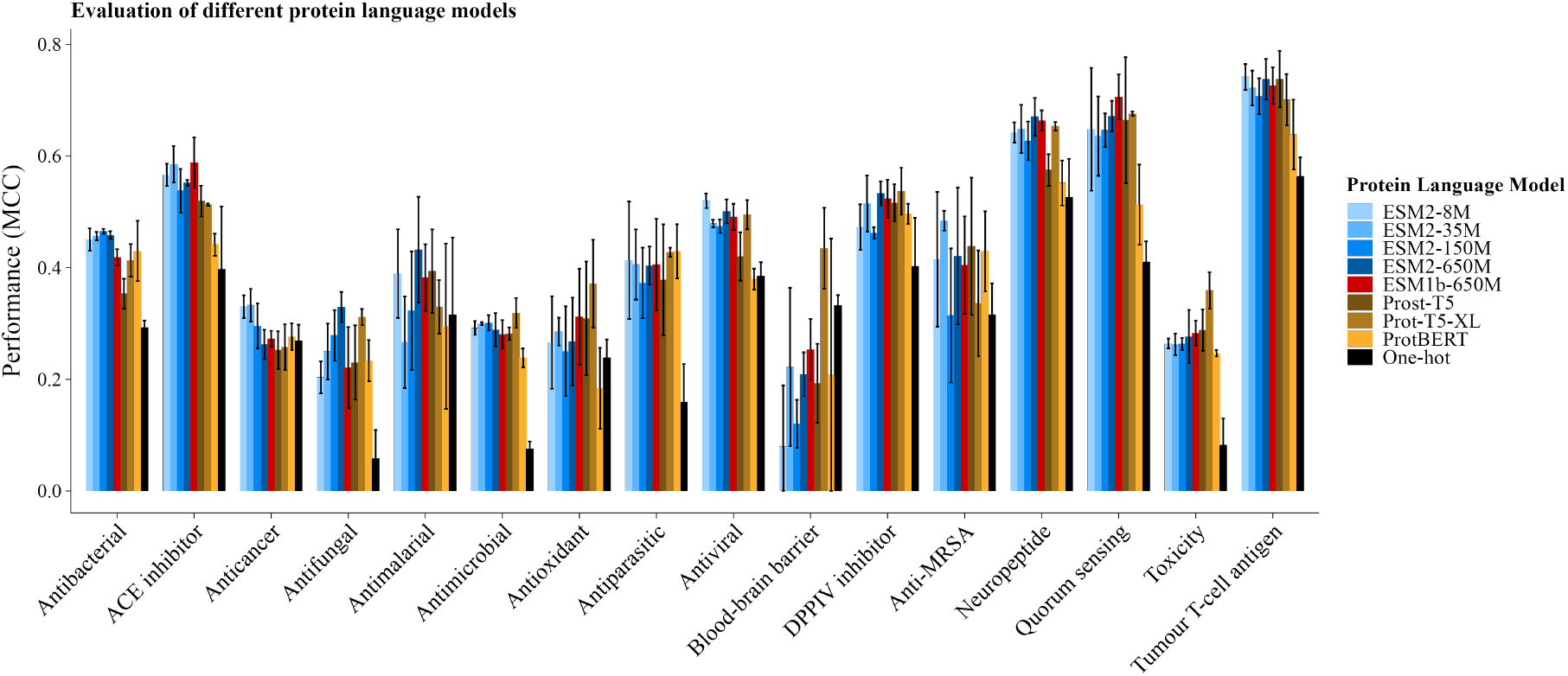
Evaluation of different protein language models. Error bars reflect the standard deviation across three replicates.

#### Model size

We evaluated four different PLM models from the ESM family with increasing size: ESM2-8M (8 million parameters), ESM2-35M (35 million parameters), ESM2-150M (150 million parameters), ESM2-650M (650 million parameters). We also evaluated ESM1b-650M (650 million parameters), from a previous version of ESM. Figure 3 shows that there is no significant difference between models across all datasets and no correlation between model size and performance.

This observation appears contrary to the established consensus that bigger PLMs tend to perform better (31; 32; 44). It is important to note that those studies focused on a very particular use of the PLMs known as full-model transfer learning. However, in our experiments, we relied on representation transfer instead. The main difference between both regimes is that in full-model transfer learning every parameter in the model is adjusted (fine-tuned) for the downstream task, whereas in representation transfer, the internal parameters of the model do not change. There are two main consequences that derive from this distinction.

First, full-model fine-tuning tuning requires the model to run several times through the training data to iteratively optimise its internal parameters. This is a computationally intensive operation, and the cost increases with model size. Representation transfer, in contrast, only requires a single run through the training data to compute the representations and is thus much faster and does not require specialised hardware like GPUs. Furthermore, when working with small datasets sizes (like most peptide datasets), there is less risk of overfitting with representation transfer (only the parameters of the downstream model are optimised) than with full-model fine-tuning (where both the model parameters, 8 − 650 × 10^6^, and the downstream models are optimised). We decided to focus on representation transfer for this study because of these two reasons.

Second, the more parameters a model has, the greater its learning capacity will be. This learning capacity can only be accessed in full-model fine-tuning as it allows for the optimisation of the internal parameters of the model. Thus, observing no correlation between model size and downstream performance in a representation transfer setting is not necessarily inconsistent with the prior literature. The question of whether the established PLM scaling rules for full-model transfer learning when modelling peptide sequences remains unanswered and is left for future work.

#### Type of model

We further compared the ESM models to the main models from the RostLab family: ProtBERT, Prot-T5-XL-UniRef50, and Prost-T5. Figure 3 shows that even though for certain datasets there might be significantly better models, when the effect is analysed across all datasets there is no significant difference between the different models or families.

All things considered, the ESM2-8M model achieves a commendable balance between enhanced performance relative to one-hot encoding and minimal computational requirements.

### Hyperparameter optimisation and ensembling of simple ML models as an alternative to neural networks

We compared the performance of two general purpose frameworks: a deep learning based model (UniDL4BioPep-A (36) and Table SD) and our optimised machine learning ensemble (OMLE).

#### Comparison to handcrafted models

Figure 4.A shows that when applied to a literature derived benchmark set of datasets, the two general purpose PLM-enabled bioactivity predictors have a performance comparable with the self-reported performance of the best handcrafted models for each specific dataset (see Table SF for the reference of each of the models). Moreover, our proposed AutoML solution (OMLE) was able to out-perform the handcrafted models on 6 out of 17 benchmark datasets for which data was available and was not significantly different for another 2. Notably, the automatic solution does not require technical expertise for development, whereas each of the handcrafted models are the result of extensive studies on peptide representation engineering and model optimisation.

**Fig. 4.**
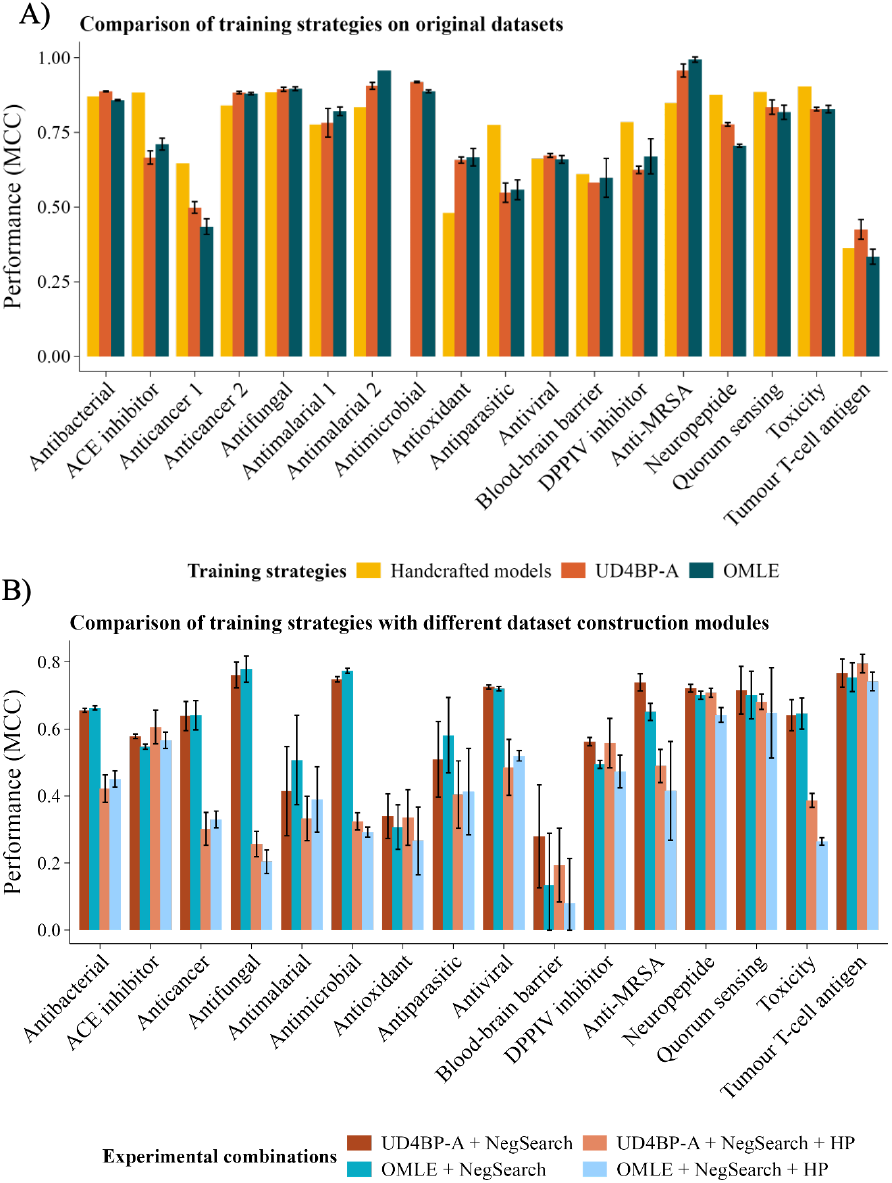
A: **Comparison of training strategies on original datasets**. B: **Comparison of training strategies with different dataset construction modules**. Error bars reflect the standard deviation across three replicates. OMLE: Optimised ML ensemble; UD4BP-A: UniDL4BioPep-A

#### Comparison of an optimised ML ensemble with a neural network

When compared with Fig 4.A, Figure 4.B shows that when the new sampling strategy for gathering negative peptides is introduced, both ML and neural network general purpose models show an equivalent drop in apparent performance, reflecting the more challenging task of predicting a specific bioactivity against peptides from a diverse collection of bioactivities. The performance drops further in both models when homology-based partitioning is introduced. Remarkably, there is no significant evidence of greater overfitting on the part of the DL model, despite the small dataset sizes, this might be due to the relatively small size of UniDL4BioPep-A. Overall, these results allow us to conclude that OMLE achieves comparable performance to a more complex neural network model, while being both more user-friendly and computationally efficient.

### AutoPeptideML

All the findings described thus far, were used to guide the development of AutoPeptideML, a computational tool and webserver that allows researchers to easily build strong peptide bioactivity predictors and provide a robust evaluation that complies with community guidelines (see Supplementary E for a more through description of the AutoML system). Figure 5 provides an overview of the final AutoPeptideML workflow.

**Fig. 5.**
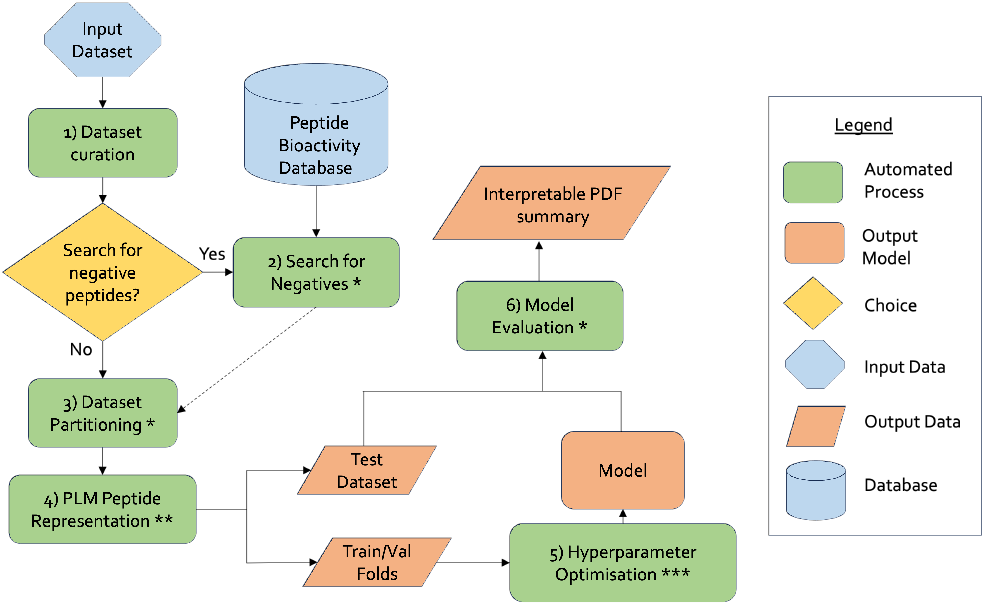
Visual summary of the AutoPeptideML workflow. *: Steps based on the results showcased in Figure 2. **: Step based on the results showcased in Figure 3. ***: Step based on the results showcased in Figure 4.

AutoPeptideML can be used in two regimes: *Model builder* and *Prediction*. In the first mode new predictive models are created automatically from a single file with known positive peptides for the bioactivity of interest. In the second mode, any predictive model generated through the *model builder* can then be used to predict for each peptide in a dataset the likelihood that it possesses the desired bioactivity.

The outputs that the program generates are:

- *Model builder:* When used to develop new predictors, AutoPeptideML outputs a model fitted to predict the bioactivity of interest, a folder with all information necessary for reproducing the model, and an interpretable summary of the model capabilities.
- *Prediction:* AutoPeptideML can also be used to leverage existing predictors. In this case, it outputs a list of the problem peptides sorted in descending order of predicted bioactivity (higher bioactivity first) and a measure of the uncertainty of each prediction.

AutoPeptideML is a tool that enables teams of experimental researchers without access to modelling expertise to quickly and easily build and interpret custom models to integrate into their experimental workflows. It can also be helpful for computational researchers to generate quick and robust baselines at the early stages of a new modelling project against which to compare any new methods. Moreover, AutoPeptideML allows for the combination of PLM representations with handcrafted features. Thus, it can assist both domain experts and computational scientists by providing a flexible and easy-to-use end-to-end modelling pipeline, allowing the former to easily construct new models and the latter to quickly run experiments and compare between different representation methods or modelling algorithms within a robust and reproducible environment.

## Conclusions

The definition of the negative class used for building peptide bioactivity predictors has a significant impact on the model performance of up to 40% and has to be controlled in order to properly interpret model predictions. Here we introduced a negative sampling strategy that gathers negative peptides from a collection of peptides with diverse bioactivities which has been shown to achieve a balance between the strengths and weaknesses of current methods in terms of the specificity and interpretability of the predictions, and the reliability of the estimation of future model performance.

The partitioning strategy used to generate training and testing subsets impacts the evaluation of model generalisation significantly and the introduction of homology-based partitioning algorithms can lead to a drop in perceived model performance of up to 50% when compared to random partitioning. The magnitude of these effects suggests that the model performance has been overestimated in most previous studies.

Using protein language models (PLMs) for computing peptide representations is a significantly better strategy than using a one-hot encoding (a naive representation) for most of the datasets considered. This underscores the potential of PLMs to compute peptide representations, in line with previous studies. Surprisingly, there is no significant correlation between model size and the performance observed, nor among different models. This marks a first step towards understanding the limitations of PLM scaling rules as it pertains their use for modelling peptide sequences.

The combination of PLM peptide representations and an optimised ensemble of simple ML models reaches state-of-the-art performance when compared both to an alternative general-purpose-framework and dataset-specific, handcrafted models across a set of 18 different datasets. Furthermore, there is no significant difference between using an ensemble of simple ML algorithms and more complex DL algorithms (UniDL4BioPep-A), even though the former is more computationally efficient.

Finally, we present AutoPeptideML as a computational tool and webserver that researchers without technical expertise to develop predictive models for any custom peptide bioactivity and facilitates compliance with community guidelines for predictive modelling in the life-sciences. It is able to handle several key steps in the peptide bioactivity predictor development life-cycle including: 1) data gathering, 2) homology-based dataset partitioning, 3) model selection and hyperparameter optimisation, 4) robust evaluation, and 5) prediction of new samples. Further, the output is generated in the form of a PDF summary easily interpretable by researchers not specialised in ML; alongside a directory that ensures reproducibility by containing all necessary information for re-using and re-training the models.

The foundational principles underlying the issues described and solutions implemented throughout this study are relevant for the application of trustworthy ML predictors for any other biosequence (e.g., DNA, RNA, proteins, peptides, DNA methylation, etc.) and their automation facilitates the rigorous evaluation and development of new models by researchers not specialised in ML.

## Acknowledgements

The authors thank Silvia González López for designing the AutoPeptideML logo, Marcos Martínez Galindo for his technical advice on how to to deploy the AutoPeptideML webserver, and Patrick Timmons for his insightful comments on an early version of the manuscript.

## A. APML-Peptipedia

The original Peptipedia database integrates information from 30 peptide bioactivity databases collecting almost 97,331 bioactive peptides labelled with 128 bioactivities (version 29_03_2023). APML-Peptipedia is the result of removing all sequences with non-standard residues or without any known bioactivity and contains 92,092 peptides (see Supplementary). Figure 1 describes the distribution of the lengths of the peptides comprising APML-Peptipedia.

**Fig. 1.**
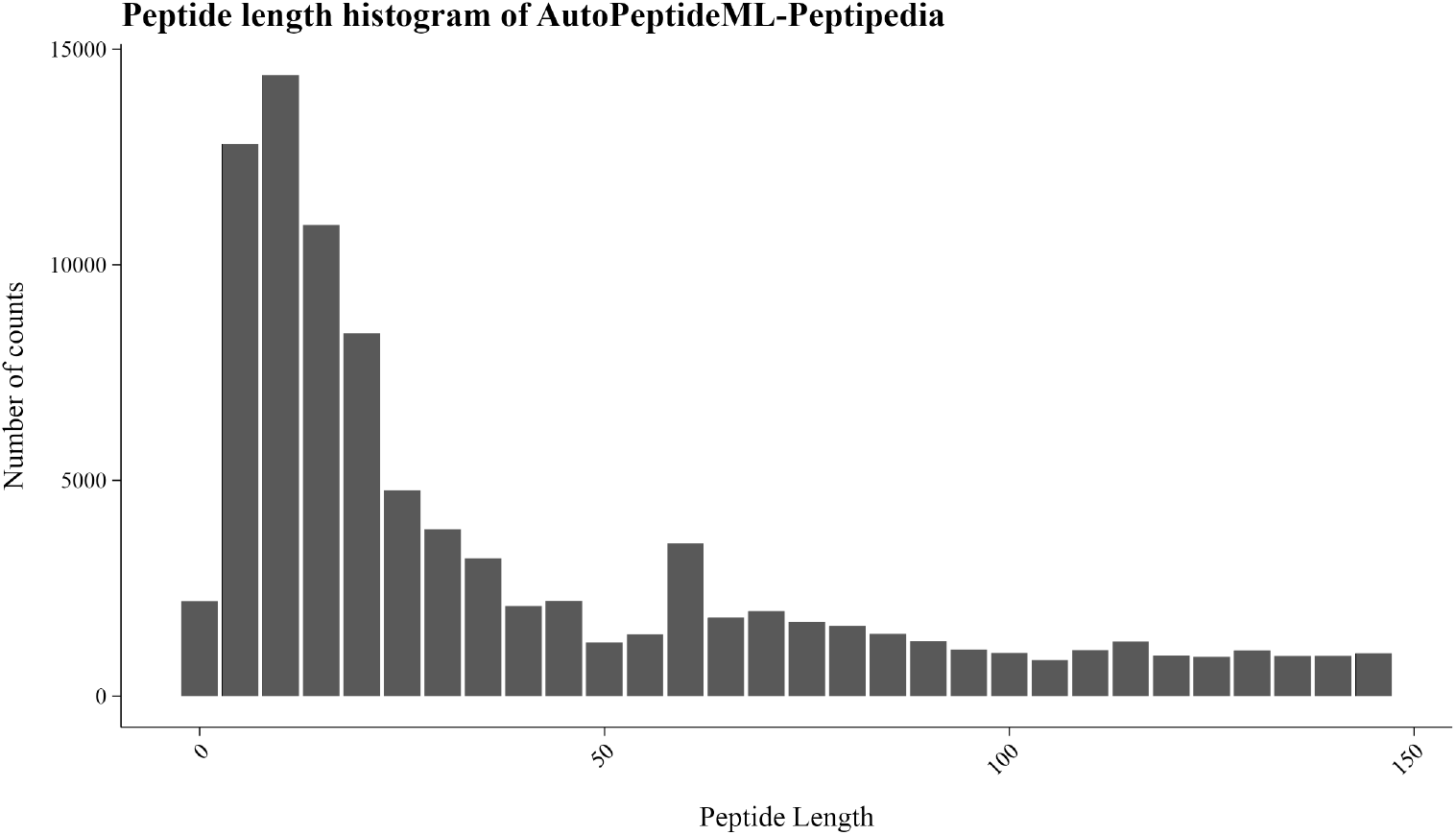
Histogram of peptide lengths with bin size 5.

## B. Search for Negative Peptides

Table 1 compiles the bioactivity tags excluded from the negative set when building the “NegSearch” datasets. The meaning behind these tags can be further expanded in the original publication (7).

**Table 1.**
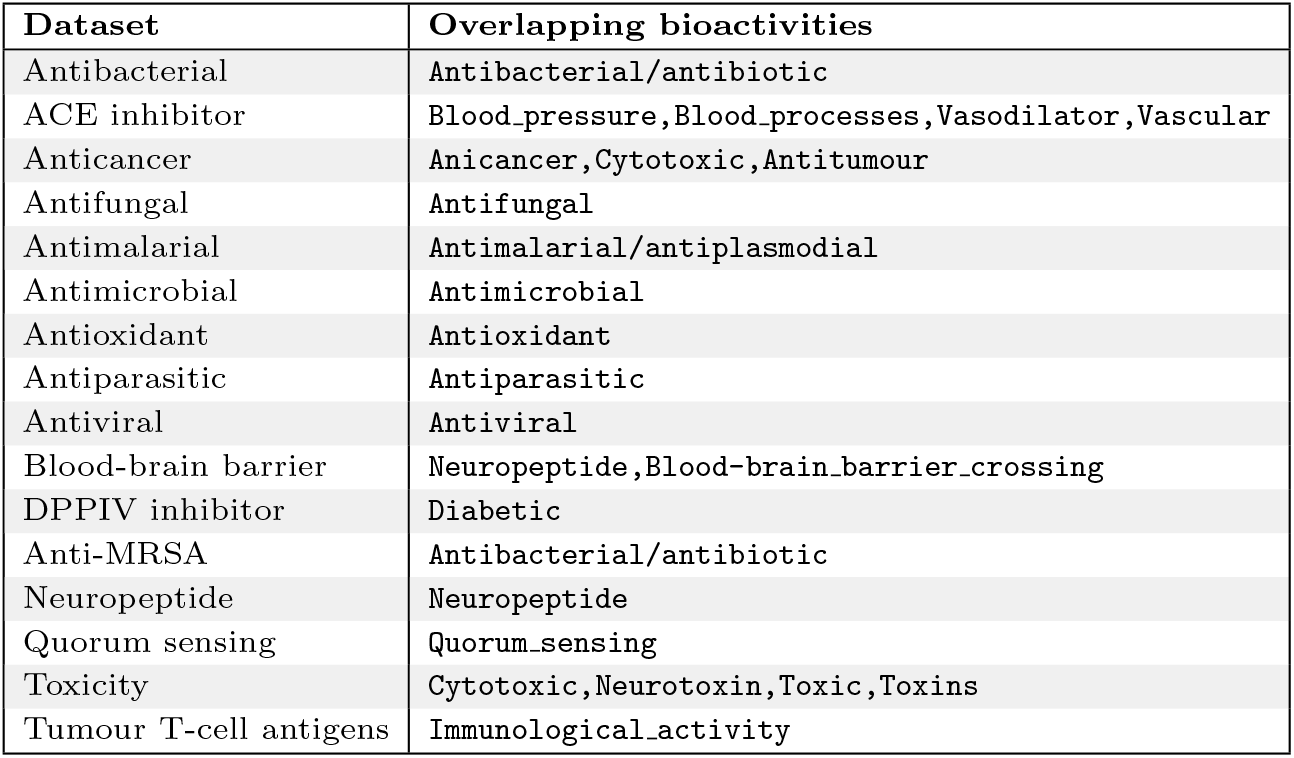
Overlapping classes excluded from the negative set for each of the benchmark datasets.

## C. Default hyperparameter search space

Table 2 describes the hyperparameter space defined for all experiments using the “Optimised ML ensemble”.

**Table 2.** Default hyperparameter search space for the ensemble used throughout the paper.

| Model | Trials | Hyperparameter search space |  |  |  |
| --- | --- | --- | --- | --- | --- |
|  |  | Name | Type | Range | Log-scale |
| KNN | 10 | K<br>Weights | integer<br>categorical | 1-30<br>uniform or distance | No<br>No |
| RFC | 10 | Max depth | integer | 2-20 | No |
|  |  | Number of estimators | integer | 10-100 | No |
| LightGBM | 10 | Max depth | integer | 1-30 | Yes |
|  |  | Number of leaves | integer | 5-50 | Yes |
|  |  | Learning rate | float | 10e-3 - 0.3 | Yes |

## D. UniDL4BioPep model architecture

UniDL4BioPep (36) computes the peptide-level representations using ESM2-8M in the same way as described in Methods, by averaging across all residue-level representations. It then uses a 1D-convolutional neural network (1D-CNN) to make the predictions. The architecture for this 1D-CNN are fixed and are described in Table 3.

**Table 3.** UniDL4BioPep architecture.

| Layer | Type of layer | Input | Size | Output |
| --- | --- | --- | --- | --- |
| 1 | Conv1D | 320 | 32 | $32 \times 320$ |
| 2 | MaxPool | $32 \times 320$ | ∼ | $32 \times 160$ |
| 3 | Flatten | $32 \times 160$ | ∼ | 5,120 |
| 4 | Dense | 5,120 | 64 | 64 |
| 5 | Output | 64 | 2 | 2 |

## E. AutoPeptideML

### Algorithm

The primary objective behind the design of AutoPeptideML is to provide a user-friendly tool that does not require extensive technical knowledge to use, while still remaining highly versatile. This is achieved through a pipeline that guarantees compliance with community guidelines such as DOME (Data, Optimisation, Model, and Evaluation) (16), ensuring a robust scientific validation (see below).

Users are free to define the number of models that should be included in the hyperparameter optimization, as well as their hyperparameter search space. AutoPeptideML supports the following algorithms: K-nearest neighbours (KNN), light gradient boosting (LightGBM), support vector machine (SVM), random forest classification (RFC), extreme gradient boosting (XGBoost), simple neural networks like the multi-layer perceptron (MLP), and 1D-convolutional neural networks (1D-CNN). Model selection and HPO are conducted simultaneously in a cross-validation regime so that the metric to optimise is the average across *n* folds. Thus, the system is never exposed to the testing set, which is kept unseen until the final model evaluation (16).

### Recommendations for using AutoPeptideML and reporting its results

This section explores how the structure of the outputs from AutoPeptideML facilitates compliance with DOME guidelines (16), nevertheless, it is important to note that no system can fully avoid its misuse or abuse and the ultimate responsibility of following proper guidelines and accurately reporting the results lies in the final users.

- **Data:** The algorithm ensures independence between the optimisation (training) and evaluation (test) sets. The hyperparameter optimisation and model selection, which can be considered as meta-optimisation strategies, relies on *n*-fold cross-validation and maintains the independence of the testing set. Further, the constraints upon the algorithm in the web-server application impedes malpractices like the manual curation of parameters to meta-optimise the results in the independent test sets. The datasets generated during the automatic search for negative samples, the train/test partitions, and the *n* train/validation folds are included in the ZIP-compressed output file, thus making their release and sharing easy. The automatic search for negatives is also compliant with the recommendation that the distribution of the data is representative of the domain in which the model is going to be applied. The use of random seeds for any stochastic process improves the reproducibility when the same exact datasets are used, thus guaranteeing that different runs will produce similar results.
- **Optimisation:** Metrics for each fold in cross-validation are provided alongside the final evaluation metrics of the model so that train versus test error can be calculated as a measure of possible under- or over-fitting. The hyper-parameter configurations of the final models are included in the output file and are therefore easy to share.
- **Model:** PLMs are not directly explainable and it follows that models built on top of their representations are thus not explainable.
- **Evaluation:** Models are evaluated with a wide array of metrics and a PDF summary of the main model performance plots and evaluation metrics is provided with a guide on how to interpret them depending on different application contexts meant for researchers that are not familiar with ML concepts. Most common problems when analysing evaluation metrics arise when working with imbalanced testing datasets, the automatic dataset construction module bypasses this problem by generating balanced datasets.

## F. Review of original benchmarks

Table 4 contains a review of the datasets used as benchmarks throughout the study: their origin, how the negatives peptides were drawn, training-testing partitioning strategy, and the reference of the best handcrafted model reported.

**Table 4.**
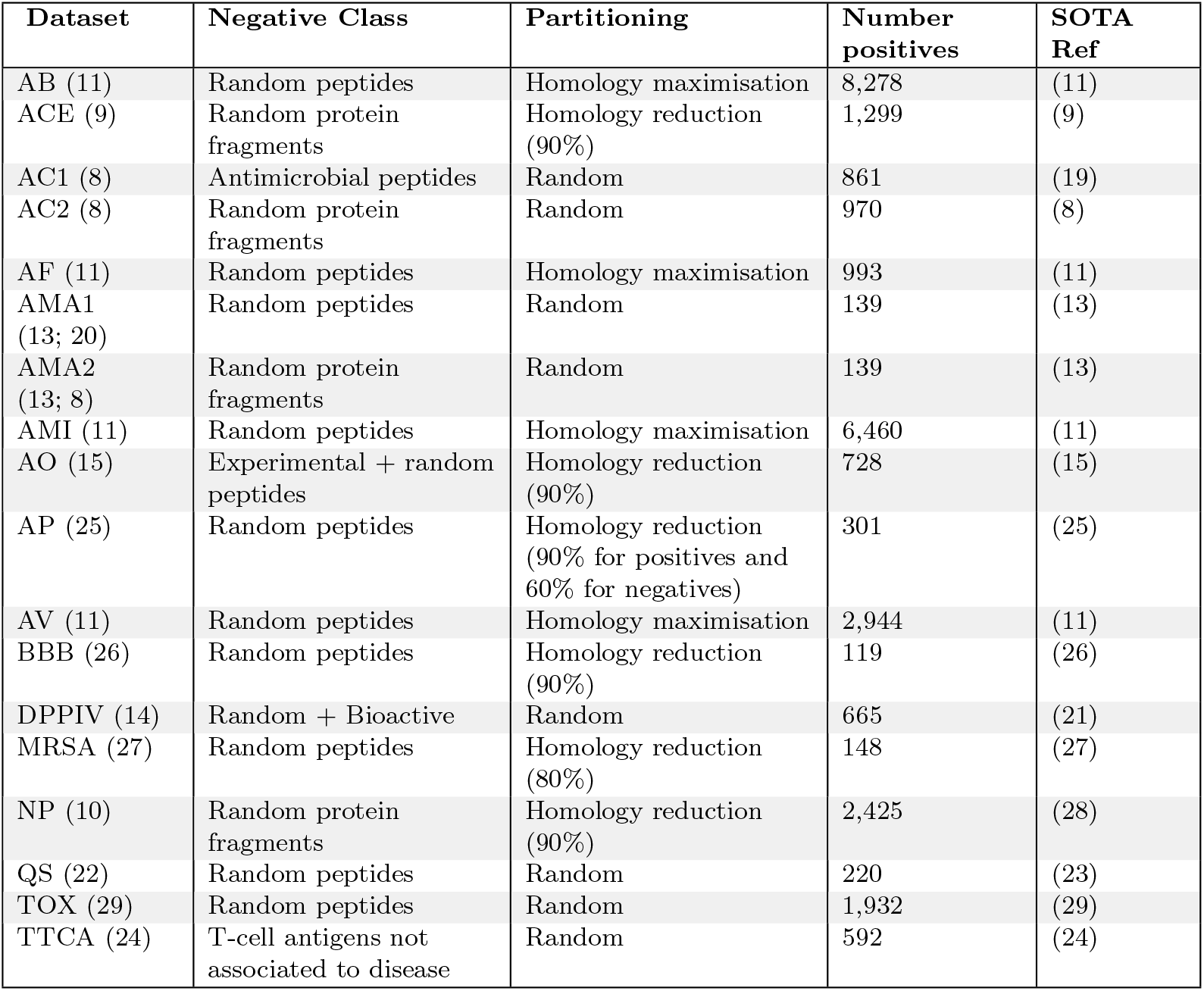
Original benchmark datasets. AB: Antibacterial; ACE: ACE inhibitor; AC: Anticancer; AF: Antifungal; AMA: Antimalarial; AMI: Antimicrobial; AO: Antioxidant; AP: Antiparasitic; AV: Antiviral; BBB: Brain-blood barrier crossing; DPPIV: DPPIV inhibitor; MRSA: Anti-MRSA; NP: Neuropeptide; QS: Quorum sensing; TOX: Toxic; TTCA: Tumor T-cell antigens; SOTA Ref: Reference to best reported model in the literature.

